## Supplemental Information for "Genome-Wide CRISPR Screen Identifies Non-Canonical NF-κB Signaling as a Potent Regulator of Density-dependent Proliferation"

SUPPLEMENTARY INFORMATION

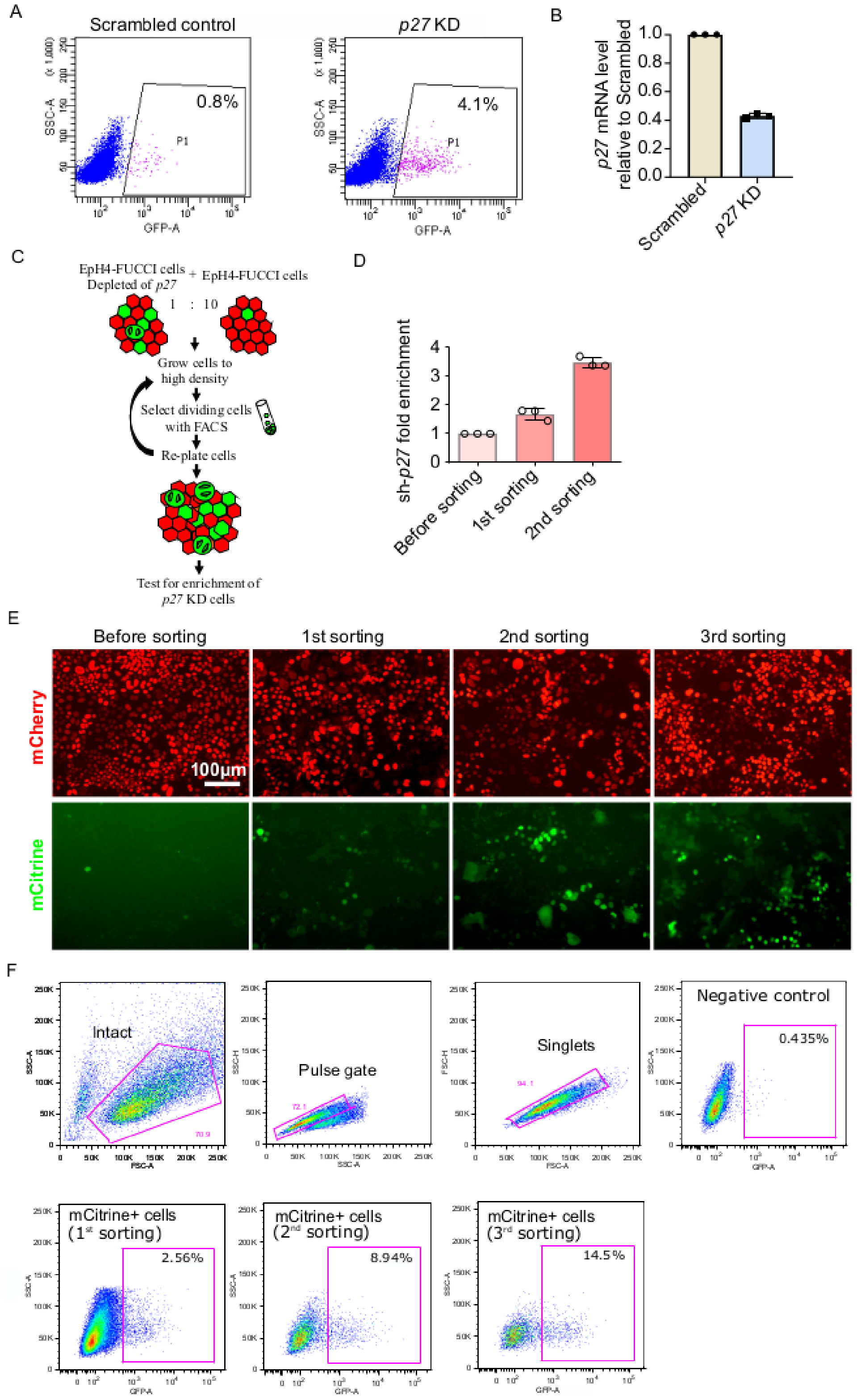

**Figure 1 - figure supplement 1.** (A) Sorting of EpH4-FUCCI cells by FACS. The number of mCitrine+ cells is increased by depletion of *p27*. (B) *p27* mRNA level in *p27* KD cells relative to scrambled control cells measured by qPCR. (C) Strategy for proof-of-principle experiments. (D) Enrichment of sh-*p27* after 1$^{st}$ and 2$^{nd}$ round of sorting, compared to sh-*p27* content before FACS, measured by qPCR. Histogram shows mean ± 1 s.d. (n=3 technical repeats) (E) Imaging of EpH4-FUCCI cells at 4 DPC before sorting and after different rounds of sorting. (F) Gating for sorting EpH4-FUCCI cells by FACS. The number of mCitrine+ cells increases after each round of sorting.

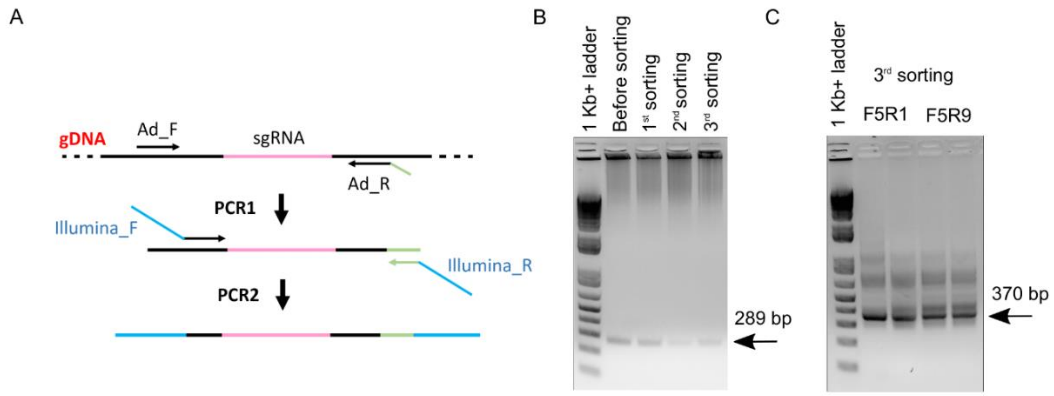


D

| Time point | Total sgRNAs | Zero counts | Gini Index |
|---|---|---|---|
| Unsorted control | 125,793 | 2808 | 0.15 |
| 1st sorting | 125,793 | 14779 | 0.39 |
| 2nd sorting | 125,793 | 10365 | 0.33 |
| 3rd sorting | 125,793 | 13612 | 0.36 |

D'

| Time Point | Reads | Mapped | Percentage |
|---|---|---|---|
| Unsorted control | 49,384,395 | 35,363,444 | 0.72 |
| 1st sorting | 21,736,214 | 13,145,089 | 0.6 |
| 2nd sorting | 21,400,849 | 15,870,963 | 0.74 |
| 3rd sorting | 71,162,949 | 51,610,375 | 0.73 |

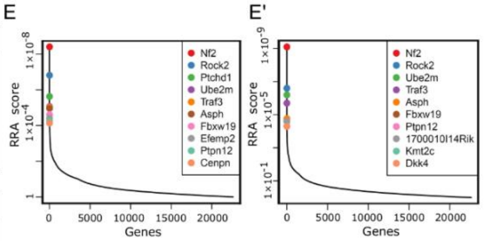


F

| Rank | ID | FDR | Enriched/total sgRNAs |
|---|---|---|---|
| 1 | *Nf2* | 0.00495 | 6/6 |
| 2 | *Rock2* | 0.01733 | 5/6 |
| 3 | *Ptchdl* | 0.09076 | 2/6 |
| 4 | *Ube2m* | 0.18069 | 5/6 |
| 5 | *Traf3* | 0.18069 | 4/6 |
| 6 | *Asph* | 0.18069 | 3/6 |

F'

| Rank | ID | FDR | Enriched/total sgRNAs |
|---|---|---|---|
| 1 | *Nf2* | 0.00495 | 6/6 |
| 2 | *Rock2* | 0.02723 | 5/6 |
| 3 | *Ube2m* | 0.02805 | 5/6 |
| 4 | *Traf3* | 0.05817 | 4/6 |

**Figure 1 - figure supplement 2.** sgRNA sequence PCR for NGS and sequence processing. (A) Schematic of PCR reactions aimed to amplify sgRNA containing fragments from gDNA (PCR1) and attach Illumina primers (PCR2) for NGS. (B) The agarose gel with DNA amplified from gDNA of control samples and samples after different rounds of sorting (PCR1). (C) An agarose gel with PCR2 products for the 3rd sorting sample with different Illumina primer pairs shown as an example of PCR2 reaction products. (D) The number of lost sgRNA (zero counts) and Gini index in samples before sorting and after different rounds of sorting. (D') Total number of reads, number of mapped reads and the fraction of mapped reads in samples before sorting and after different rounds of sorting. (E), (E') RRA score plots for samples after 1st and 2nd sorts, respectively. (F), (F') The list of genes with FDR below 0.25 and more or equal to three sgRNAs enriched compared to control after 1st and 2nd sort, respectively.

**Figure 1 - figure supplement 3.** Control shRNA and qPCR primers sequences used in this study.

| | |
|---|---|
| control shRNA | CCGGTCCTAAGGTTAAGTCGCCCTCGCTCG AGCGAGGGCGACTTAACCTTAGGTTTTTG |
| *p27*-qPCR-FWD | TCAAACGTGAGAGTGTCTAACG |
| *p27*-qPCR-REV | CCGGGCCGAAGAGATTTCTG |
| Puromycin-qPCR-FWD | CTGCAAGAACTCTTCCTCACG |
| Puromycin-qPCR-REV | GGGAACCGCTCAACTCGG |

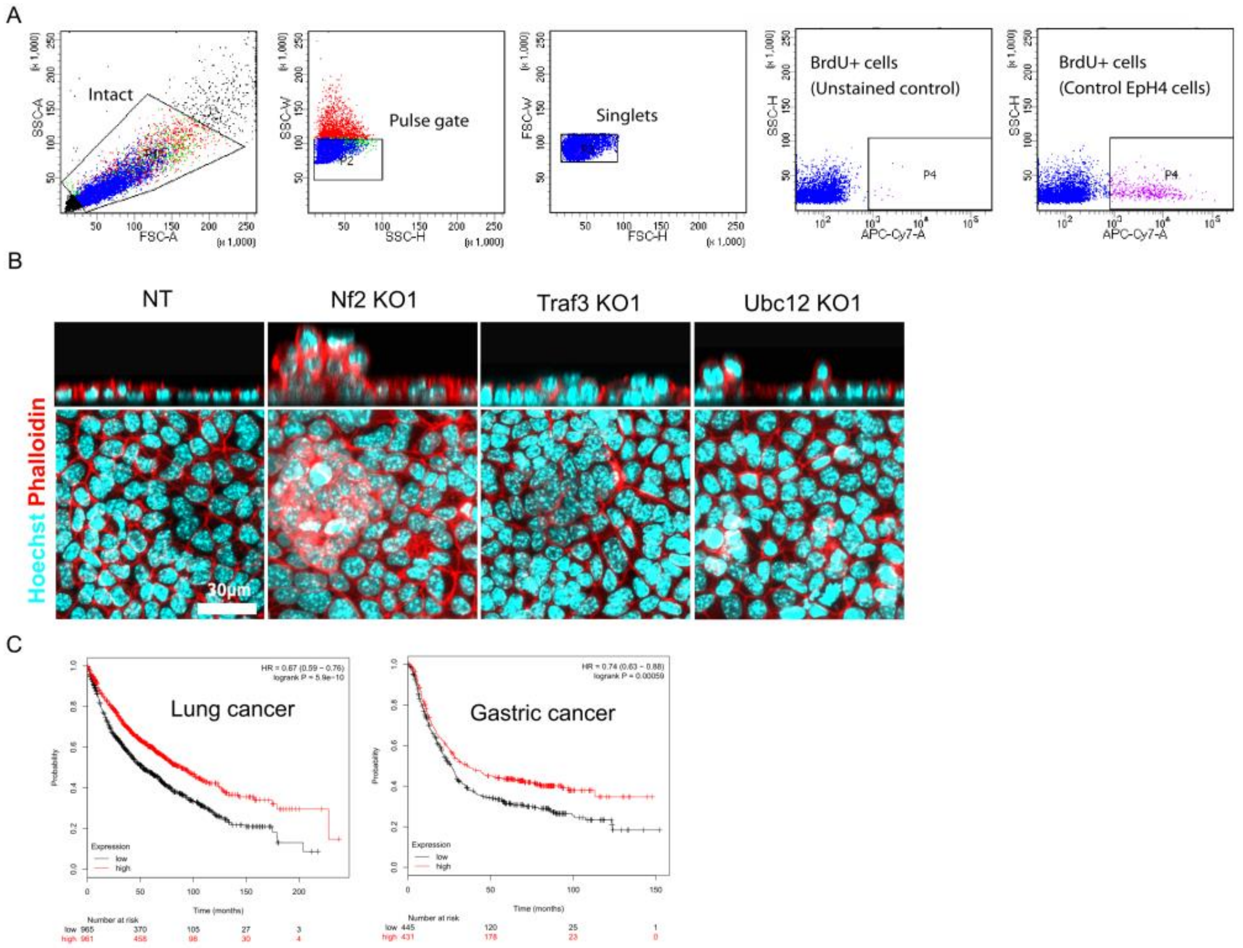


**Figure 2 - figure supplement 1.** BrdU+ cell cytometry gates and cancer survival based on *Traf3* expression level. (A) Flow cytometry gating for cells stained for BrdU. (B) Control and KO cells stained by phalloidin and Hoechst. Top panel shows xz view. (C) Kaplan-Meier plots show overall survival in all lung and gastric cancer based on *Traf3* expression level. Red and black curves reflect patients with high and low *Traf3* expression, respectively.

**Figure 2 - figure supplement 2.** sgRNA sequences used in this study.

| | |
|---|---|
| control non-targeting (NT) | GCGAGGTATTCGGCTCCGCG |
| mouse *Nf2* sgRNA KO1 | CGAGATGGAGTTCAACTGCG |
| mouse *Nf2* sgRNA KO2 | ATACTGCAGTCCAAAGAACC |
| mouse *Traf3* sgRNA KO1 | GTGCTCGTGCCGGAGCAAGG |
| mouse *Traf3* sgRNA KO2 | TGGCCCTTCAGGTCTACTGT |
| mouse *Ubc12* sgRNA KO1 | GCGCAGCTCCGGATTCAGAA |
| mouse *Ubc12* sgRNA KO2 | GAGTCGGCCGGCGGCACCAA |
| mouse *NF-κB2 (p100)* sgRNA KO1 | CTGAGCGTGATAAATGACGT |
| mouse *NF-κB2 (p100)* sgRNA KO2 | CTGTTCCACAATCACCAGAT |
| mouse *Map3K14 (Nik)* sgRNA KO1 | TCAGAGCGCATTTTCATCGC |
| mouse *Map3K14 (Nik)* sgRNA KO2 | GTCGAGGCAGTACCGGTCGC |
| human *TRAF3* sgRNA KO1 | AGATTCGCGACTACAAGCGG |
| human *TRAF3* sgRNA KO2 | CCTCACATGTTTGCTCTCGC |

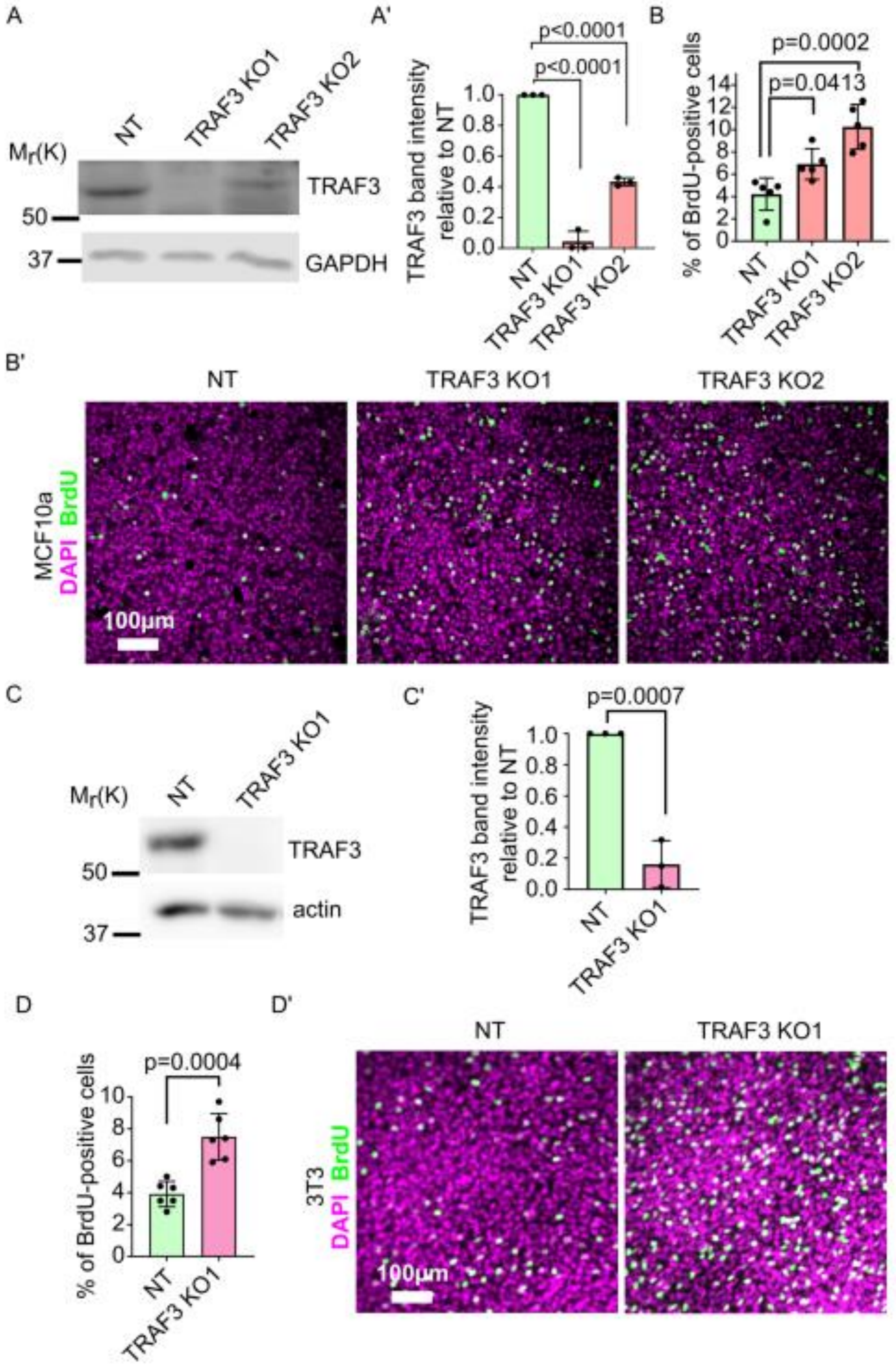


**Figure 3 - figure supplement 1.** Loss of TRAF3 in MCF10a and NIH 3T3 cells causes over-proliferation. (A) Immunoblotting of NT and TRAF3 KO MCF10a cells for TRAF3. GAPDH was used as a loading control. (A') Quantifications of TRAF3 levels based on Western blots (A)

Histogram shows mean ± 1 s.d. (n=3). P values were calculated by one-way ANOVA followed by Dunnett's multiple comparisons test. (B) Cytometric analysis of NT control and TRAF3 KO MCF10a cells stained for BrdU to assess proliferation at 4 DPC. Histogram shows mean ± 1 s.d.(n=3). P values were calculated by one-way ANOVA followed by Dunnett's multiple comparisons test. (B') NT control and TRAF3 KO MCF10a cells at 4 DPC were treated and stained for BrdU and DAPI as nuclear marker. (C) Immunoblotting of NT and TRAF3 KO NIH 3T3 cells for TRAF3. actin was used as a loading control. (C') Quantifications of TRAF3 levels based on Western blots (C) Histogram shows mean ± 1 s.d. (n=3). P values were calculated by Student t-test. (D) Cytometric analysis of NT control and TRAF3 KO NIH 3T3 cells stained for BrdU to assess proliferation at 4 DPC. Histogram shows mean ± 1 s.d. (n=3). P values were calculated by Student t-test. (D') NT control and TRAF3 KO NIH 3T3 cells at 4 DPC were treated and stained for BrdU and DAPI as nuclear marker.

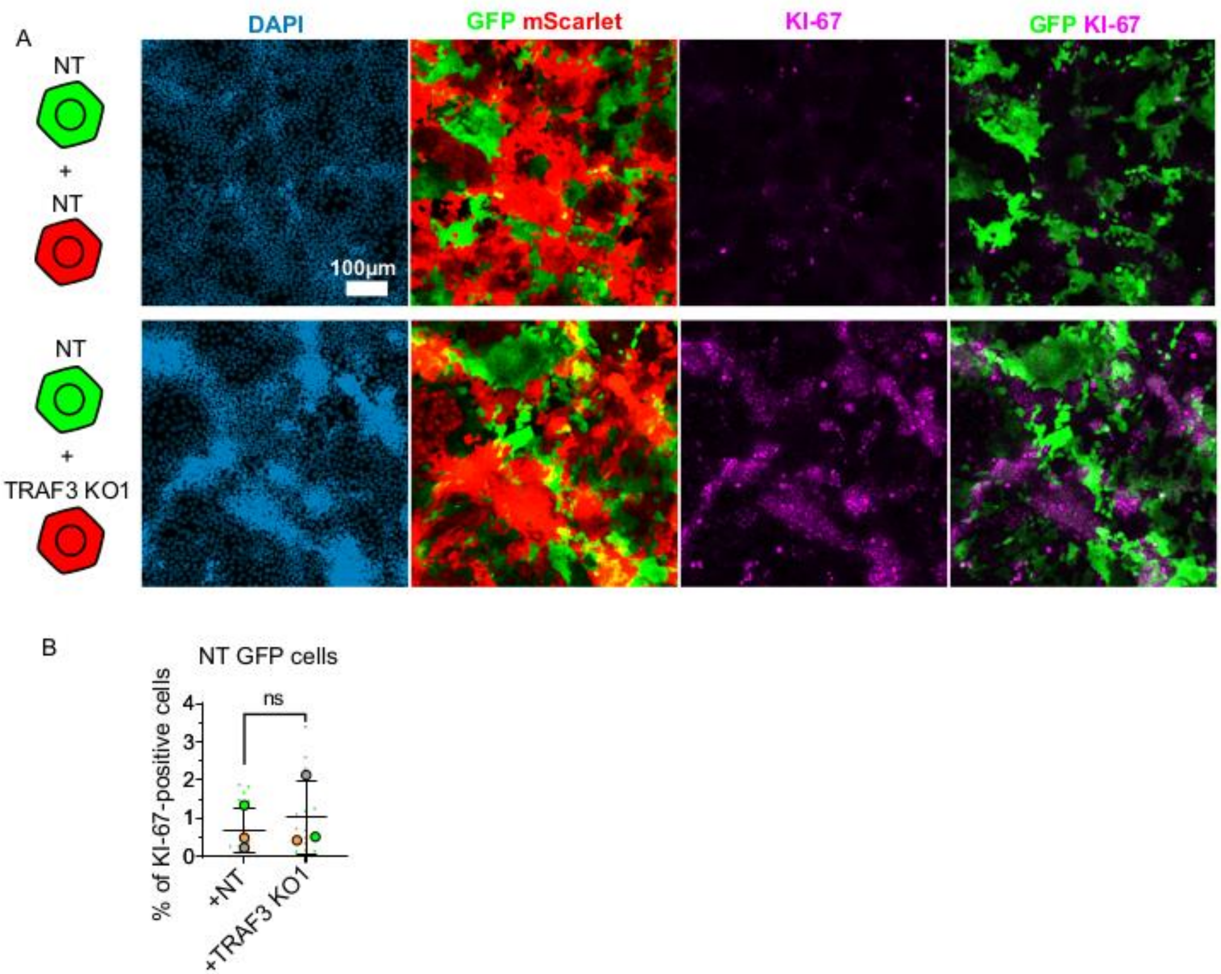


**Figure 7 - figure supplement 1.** TRAF3 KO cells over-proliferate cell autonomously. (A) NT and TRAF3 KO1 cells labeled with GFP or mScarlet were mixed at a 1:1 ratio (NT GFP + NT mScarlet, NT GFP + TRAF3 KO1 mScarlet), and grown to 4 DPC. Cells were stained for KI-67 and DAPI. (B) Quantifications of NT GFP cell proliferation in mixture with NT mScarlet or TRAF3 KO1 mScarlet. Proliferation was measured as percent of KI-67-positive cells to the total number of NT GFP cells. Data are presented as a SuperPlot (n=3). P value was calculated by mixed model two-way ANOVA.

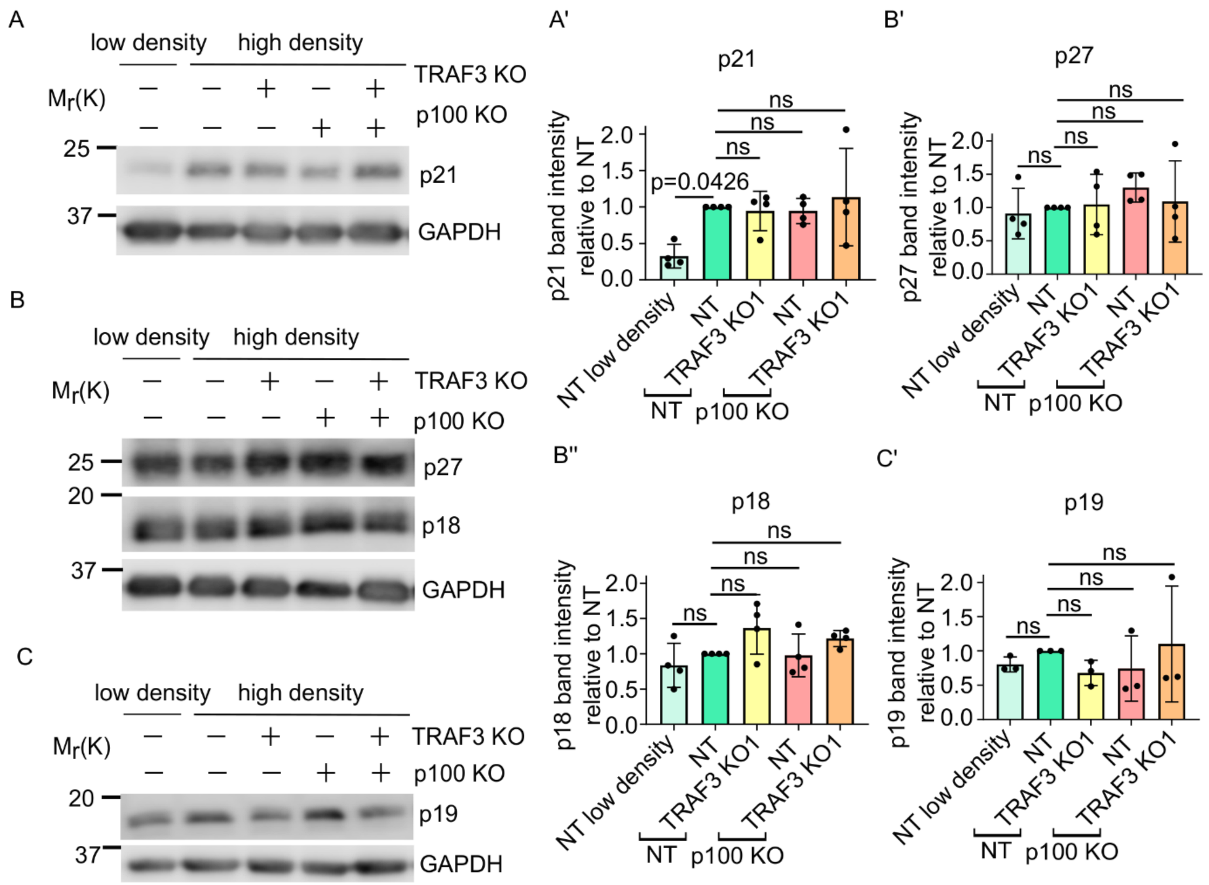


**Figure 7 - figure supplement 2.** Loss of Traf3 does not affect CKI levels. (A) Immunoblotting for p21 in sparse NT cells, dense NT, TRAF3 KO1, NT/p100 KO and TRAF3/p100 double KO cells. GAPDH was used as a loading control. (B) Western blotting for p27 and p18 in sparse NT cells, dense NT, TRAF3 KO1, NT/p100 KO and TRAF3/p100 double KO cells. GAPDH was used as a loading control. (C) Western blotting for p19 in sparse NT cells, dense NT, TRAF3 KO1, NT/p100 KO and TRAF3/p100 double KO cells. GAPDH was used as a loading control. (A')-(C') Quantifications of the blots (A)-(C). N=4 for blots (A) and (B). N=3 for blot (C) Histograms show mean ± 1 s.d. P values were calculated by one-way ANOVA followed by Dunnett's multiple comparisons test.
